# MDMA impairs response to water intake in healthy volunteers

**DOI:** 10.1101/021113

**Authors:** Matthew J. Baggott, Kathleen J. Garrison

## Abstract

Hyponatremia is a serious complication of 3,4-methylenedioxymethamphetamine (MDMA) use. We investigated potential mechanisms in two double-blind, placebo-controlled studies. In study 1, healthy drug-experienced volunteers received MDMA or placebo alone and in combination with the alpha-1 adrenergic inverse agonist prazosin, used as a positive control to release antidiuretic hormone (ADH). In study 2, volunteers received MDMA or placebo followed by standardized water intake. MDMA lowered serum sodium, but did not increase ADH or copeptin, although the control prazosin did increase ADH. Water loading reduced serum sodium more after MDMA than after placebo. There was a trend for women to have lower baseline serum sodium than men, but there were no significant interactions with drug condition. Combining studies, MDMA potentiated the ability of water to lower serum sodium. Thus, hyponatremia appears to be a significant risk when hypotonic fluids are consumed during MDMA use. Clinical trials and events where MDMA use is common should anticipate and mitigate this risk.

## Introduction

3,4- Methylenedioxymethamphetamine (MDMA, commonly referred to as ‘ecstasy’) is a widely used recreational drug. In the United States in 2013, reported lifetime/previous 30-day MDMA use was 6.8%/0.3% for persons aged 12 and older (NSDUH). Hyponatremia, defined as serum sodium concentrations of less than 135 mEq/L, is a potentially serious MDMA complication. Many published case reports document the significant morbidity and mortality of MDMA-related hyponatremia [1–20]. Symptomatic hyponatremia results from passive flow of water into cells, which can cause cerebral edema and may result in brain stem herniation. Clinical manifestations include nausea, vomiting, headache, and mental status changes. In severe cases, hyponatremia may lead to seizures, coma, and death.

Hyponatremia after MDMA use is thought to involve a combination syndrome of inappropriate antidiuretic hormone secretion (SIADH) and increased hypotonic fluid intake. However, the relative roles of ADH versus fluid intake remain unclear. ADH normally is secreted when effective circulating blood volume is decreased. SIADH is marked by increased secretion of ADH despite normal circulating blood volume, resulting in plasma hyponatremia and hypo-osmolality along with impaired free water excretion. Increased secretion of ADH has been documented in some case reports of MDMA-related hyponatremia [2, 3, 10, 12, 17, 18]. In other cases, inappropriately high urine osmolality in the setting of low serum osmolality provides additional evidence for elevated ADH [reviewed in 17]. Of note, in these case reports, there is an apparent lack of a dose-response relationship.

Controlled studies found that MDMA (and its metabolites) may induce ADH release. Henry *et al*. [21] administered 40 mg MDMA to eight males and reported an acute increase in ADH accompanied by a small decrease in serum sodium and an increase in urine osmolality [21–24]. Dumont et al. [25] administered 100 mg MDMA to 9 males and 7 females volunteers and saw serum sodium changes similar in magnitude to those seen by Henry and colleagues despite the considerable difference in dose. This suggests MDMA may not linearly increase plasma ADH. Consistent with this possibility, plasma MDMA and plasma ADH concentrations negatively correlated at 1 hour in the dataset of Henry and colleagues, which is the opposite of what would be predicted if MDMA induced SIADH. This lack of relationship may due to MDMA metabolites such as HMMA contributing to ADH release, as has been shown in rat hypothalamus *in vitro* [21–24]. A study of ecstasy users by Aitchison et al. [26] found genetic polymorphisms predicted low CYP2D6 or COMT enzyme activities were associated with greater MDMA-induced lowering of plasma sodium. This again is consistent with a role for metabolites.

In addition, the relationship between MDMA and ADH concentrations may be obscured due to difficulty assaying ADH. ADH is highly bound to platelet, which can produce artefactually high or low measurements depending on handling of samples[27–29]. Copeptin, the C-terminal part of the ADH precursor preprovasopressin, was proposed as a biological proxy for ADH because it is released as a cofactor with ADH and is reliably assayable [30]. Simmler, Hysek, and Liechti [31] reported that 125 mg MDMA elevated plasma copeptin at 60 and 120 min compared with placebo in women but not in men. Wolff et al. [32] prospectively compared dance club attendees who went on to use MDMA with those who did not and detected significant differences in plasma and urine osmolality but not significant differences in ADH when MDMA users were compared to non-users. Taken together, this literature shows modest and inconsistent effects of MDMA on ADH levels.

Alternatively, rather than directly increasing ADH, MDMA-related hyponatremia may be secondary to some other pathophysiology. This includes hypothetical effects on gastrointestinal tract motility [12] or on renal collecting tubule functioning. Drugs including fluoxetine, oxcarbazepine, and carbamazepine were reported to induce hyponatremia without a concomitant increase in ADH [33, 34]. Fluoxetine was further shown to increase expression of aquaporin 2 channels in the inner medullary collecting duct, increasing water reabsorption independent of ADH [35].

Hyponatremia after MDMA also likely has behavioral risk factors. The sudden historic appearance of MDMA-related hyponatremia strongly suggests a behavioral component [19]. MDMA-related hyponatremia was not reported until 1993, after harm reduction efforts began to recommend water consumption in an attempt to reduce risks of exercise-related dehydration and hyperthermia. In some hyponatremia case reports, witnesses reported that the individual consumed large amounts of water [4, 7, 13, 15, 18, 19, 36]. Thus, ecstasy-related hyponatremia may be partially due to erroneous user beliefs that water consumption reduces ecstasy toxicity.

There is a strong gender imbalance in MDMA-related hyponatremia. The vast majority of MDMA-related cases involve females who are less than 30 years-old and who ingested a single dose of ecstasy [2]. In contrast, most other syndromes of MDMA-related toxicity predominantly involve males. The high prevalence of females in MDMA-related hyponatremia is also seen in hyponatremia from other causes [e.g., 35, 37–39]. An elevated risk of hyponatremia symptoms in women is partly explained by the inhibitory effects of estrogen on brain Na*-KATPase, which elevates risk of cerebral edema [40]. Lower body weight and decreased muscle mass in females are also risk factors [12]. Sex differences in MDMA effects have been reported in Sprague-Dawley [41–43] and Wistar rats [44], which can at least in part be attributed to sex differences in MDMA pharmacokinetics [42]. Less is known about possible gender differences in MDMA pharmacokinetics in humans [45] and whether these contribute to the disorder.

We sought to investigate if a change in ADH is a mechanism of MDMA-induced hyponatremia in two controlled studies with healthy, MDMA-experienced volunteers. In study 1, we used the alpha-1 adrenergic inverse agonist prazosin as a positive control to stimulate ADH release by decreasing blood pressure. We sought to test whether MDMA would increase ADH and whether this would be correlated with serum sodium decreases. In study 2, we investigated the effects of water loading on indices of hydration. We hypothesized that MDMA would increase the effects of water loading on serum sodium.

### Materials and Methods

The current manuscript describes the effects of MDMA and water loading on plasma ADH and serum sodium. Additional self-report and computerized neurocognitive tasks measures relating to emotional effects of MDMA will be described in separate manuscripts.

**Participants.** We recruited healthy, MDMA-experienced individuals between the ages of 18 and 50, through newspaper and on-line advertisements and word-of-mouth. A physician determined participants to be healthy based on medical questionnaires, laboratory screenings, and a physical exam. Exclusion criteria included: DSM-IV dependence on MDMA or any other psychoactive drug (except nicotine or caffeine); desire to quit or decrease MDMA use; history of adverse reaction to study drugs; current enrollment in a drug treatment program; current supervision by the legal system; any current physical or psychiatric illness that might be complicated by the study drugs or that might impair ability to complete the study; body mass index (weight/height^2^) greater than 30 or less than 18 kg/m^2^; and current or recent use of any medication that might pose a risk of drug-drug interaction.

**Description of Procedures or Investigations Undertaken.** Both studies used a double-blind, placebo-controlled, within-subject, sequence-and gender-balanced design. We selected 1.5 mg/kg MDMA, measured as the hydrochloride salt (equivalent to 1.26 mg/kg as the freebase), as an active dose. We chose a dose that would produce typical drug effects without clinically significant changes in physiological parameters or detectible harm to participants, based on past clinical studies [e.g., 46, 47–50]. Oral administration of MDMA or its placebo took place between 10 and 10:30 am.

**Study 1 design.** In four experimental sessions that were separated by at least one week, outpatient volunteers experienced the following conditions: (a) Placebo prazosin followed one hour later by placebo MDMA; (b) Placebo prazosin followed one hour later by 1.5 mg/kg oral MDMA; (c) 1 mg oral prazosin followed one hour later by placebo MDMA; or (d) 1 mg oral prazosin followed one hour later by 1.5 mg/kg oral MDMA. The first two participants (1 male, 1 female) received two mg prazosin. However, postural hypotension, an anticipated effect of prazosin, persisted for approximately 9-11 h after MDMA. In response, we lowered the dose to 1 mg for the remaining fourteen individuals. To minimize effects of water consumption, participants were limited to drinking 1 pt (about 473 mL) or less of water each hour.

**Study 2 design.** In two experimental sessions that were separated by at least one week, participants were admitted into a hospital research ward on the evening before drug administration and fed standardized meals for dinner (including 500 mL water) and breakfast (≤ 500 mg sodium, finished at least 1.5 h before dosing). Volunteers experienced the following dosing conditions: (a) Placebo followed one hour later by oral water challenge; (b) 1.5 mg/kg oral MDMA followed one hour later by oral water challenge. Water challenge was performed by having participants drink 20mL/kg of water within 30 minutes. No other fluid intake was allowed until at least 6 h after MDMA/placebo administration.

**Materials**. MDMA was kindly provided by David Nichols, Ph.D. (Purdue University, IN). Prazosin was purchased commercially. We used opaque gelatin capsules to hold MDMA and over-encapsulate prazosin. We used lactose as placebo.

**Ethics**. We conducted this research according to the code of ethics established by the declaration of Helsinki as amended in Edinburgh. The California Pacific Medical Center Institutional Review Board approved the studies. MDMA and prazosin were administered under an Investigational New Drug exception for MDMA from the Food and Drug Administration. Volunteers provided written consent after being informed both orally and in writing of the study procedures.

**Safety monitoring**. Safety monitoring included negative urine drug screening (and, if female, negative urine test for pregnancy) immediately prior to dosing, and monitoring of vital signs until at least six h after MDMA/placebo dosing. Researchers were trained to monitor for symptoms of hyponatremia or other toxicity and were continually present for at least 8 h.

**Measures.** We collected blood samples for ADH and plasma sodium at baseline and 1, 2, 3, 4, 6 and 24 h after dosing. We collected urine in pooled samples before dosing, and 0-8 and 8-24 h after. We measured plasma sodium (mmol/L) using Siemens Dimension RxL Max Integrated Chemistry System and determined plasma ADH concentrations with Radio Immunoassay (RIA). For sodium, the inter-assay coefficients of variation were 0.74% and 1.11% respectively at 119 mg/dL. We measured plasma ADH with RIA in a clinical laboratory (Quest Diagnostics, San Juan Capistrano, California). For ADH, the lower limit of detection was 1.0 pg/mL and inter-assay and intra-assay coefficients of variation were 5.4% and 7.1% respectively. The reported reference range was 1.0-13.3 pg/mL. We measured copeptin in study 2 only using enzyme-linked immunosorbent assay (Phoenix Pharmaceuticals, Burlingame, California) at baseline and 2 and 3 h after dosing. The lower limit of detection was 32 pg/mL, and the inter-and intra-assay coefficients of variation were less than 4%. In study 1 only, we collected blood samples to measure MDMA and its metabolites (4-hydroxy-3-methoxymethamphetamine [HMMA], 4-hydroxy-3-methoxyamphetamine (HMA) and 3,4-methylenedioxyamphetamine (MDA) before and 1, 2, 4, 8, 24, and 48 h after MDMA, using the method of Scheidweiler and Huestis [51].

**Statistical and pharmacokinetic analysis**. We analyzed data using mixed-effects models in R [52] with drug condition as a fixed effect and participant as a random effect using a 2-tailed 0.05 level of significance. When analyses identified a main effect of drug condition, we made pairwise comparisons using Tukey’s HSD, and an additional model was made with gender included as a fixed effect, although this was regarded as exploratory given that the sample size limited statistical power. We checked the normality of error terms and log10-transformed data when errors were not normally distributed. Repeated measures were generally transformed to Emax [53] summary measures before analysis. Correlations were calculated using Kendall’s tau.

We estimated pharmacokinetic parameters in NONMEM using a noncompartmental model to calculate t1/2, area-under-the-curve from 0 to infinity (AUC∞), apparent volume of distribution after oral administration (Vd/F, where F is bioavailability), and apparent total body clearance of the drug from plasma (CL/F). Three or more time points were included in the calculation of t1/ 2. AUCœ was calculated by extrapolating AUC from time of dosing to infinity based on the last observed concentration and the first-order rate constant (λΖ) associated with the terminal (log-linear) portion of the curve (estimated by linear regression of time versus log concentration).

## Results

**Study 1** included sixteen (eight male and eight female) participants, ages 26.6 ± 1.8 years (mean ± SEM; range: 18-42). Males weighed 75.3 ± 4.1 kg (range: 57.3-94.1), while females weighed 63.1 ± 3.6 kg (range: 52.7-81.4). Participants drank 1213 ± 76 ml water in the 8 h following MDMA administration. Water intake did not differ by condition, body weight, or gender.

**MDMA and metabolites.** Pharmacokinetics of MDMA and metabolites are summarized in **Figure 1** and **Table 1.** There were no significant effects of gender or condition (i.e., prazosin pretreatment) on MDMA kinetics (measured as maximum concentration, area under the concentration versus time curve, or elimination half-life).

**Figure 1.**
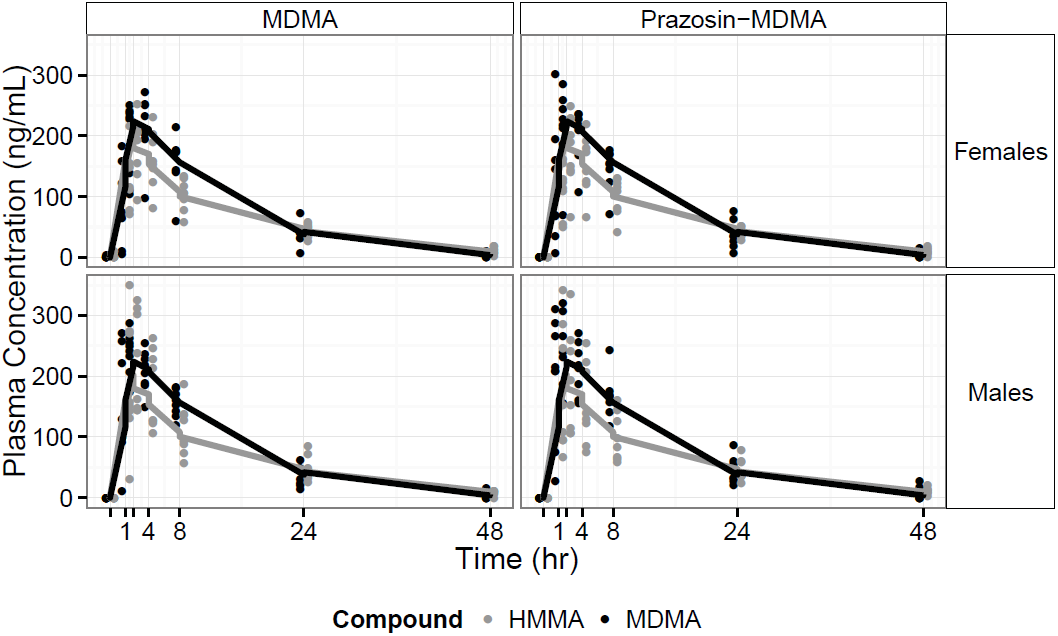
Plasma concentration versus time curves for MDMA (black) and HMMA (grey) from Study 1 after MDMA alone or prazosin with MDMA in male and female participants

**Table 1.**
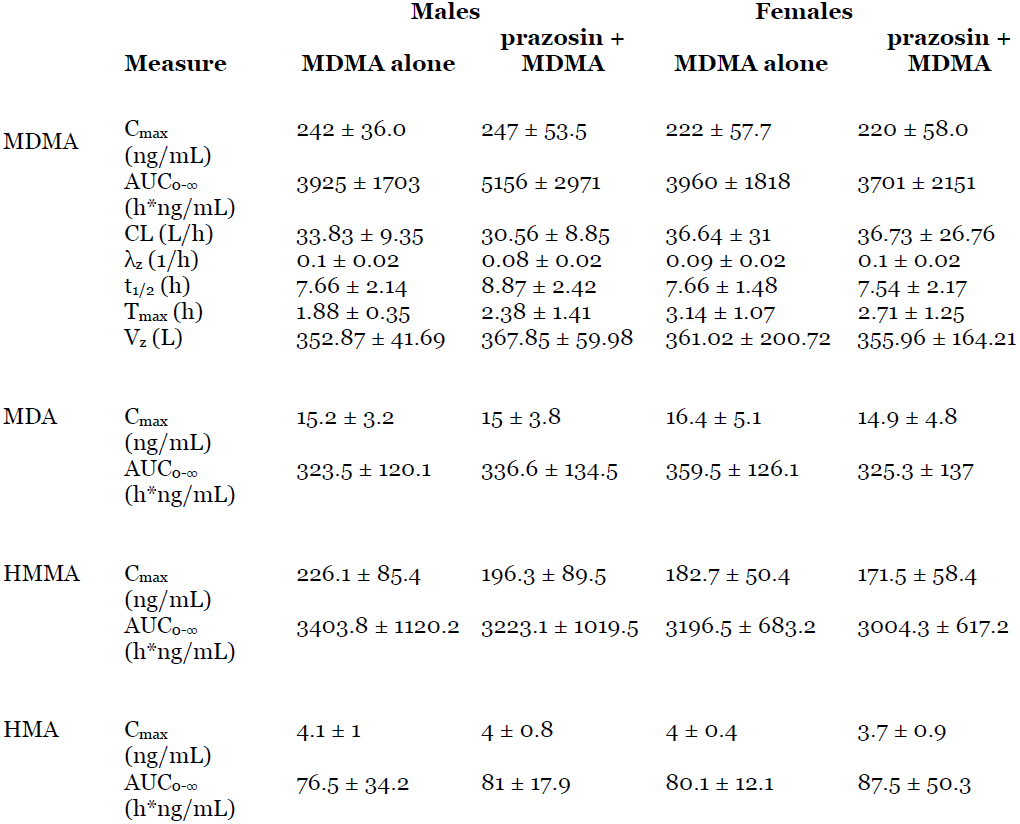
Pharmacokinetic results from Study 1. Values are given as Mean ± SD. Abbreviations: AUC0-∞ area under the curve from 0 to extrapolated to infinity, CL/F plasma clearance, Cmax maximum plasma concentration, tmax time of maximum plasma concentration, Vz/F apparent volume of distribution, Xz first order elimination constant, t_1/2_ half-life.

**ADH.** Prazosin but not MDMA increased ADH (**Figure 2**). In a model predicting maximum log10-transformed serum ADH levels with condition and gender fixed effects and participant as a random effect, there was a significant effect of condition on ADH (F_3,45_ = 5.68, p = 0.002) but not gender (p = 0.11). Both prazosin conditions increased ADH compared to placebo (Prazosin alone: z = 3.46, p = 0.002; Prazosin with MDMA: z = 3.22, p = 0.004). While a few (male) participants appeared to have elevations in ADH at 1 h after MDMA, this condition was not significantly different from placebo in our model (z = 1.21, p = 0.589). Eight of 448 ADH samples were either not collected or not analyzable.

**Figure 2.**
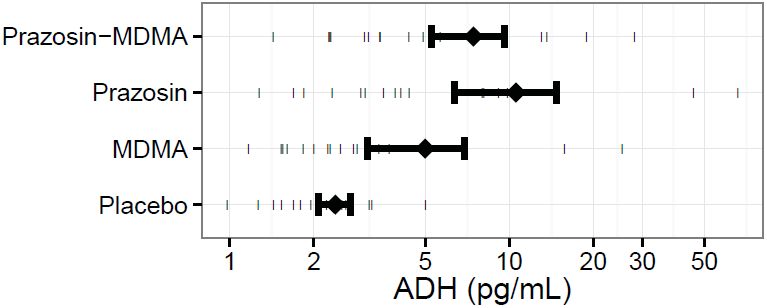
Antidiuretic hormone (ADH) after placebo, MDMA, prazosin, and prazosin with MDMA in Study 1. Diamonds and Error bars indicate mean and SEM, respectively.

**Serum sodium.** Both MDMA conditions decreased serum sodium (**Figure 3, top**). In a model predicting minimum serum sodium concentration using condition and gender as fixed effects and participant as a random effect, there were effects of condition (F_3,357_ = 8.44, p < 0.0001) and gender (F_1,14_ = 5.913, p = 0.0291) but no interaction. MDMA alone (t_357_ = -4.05, p = 0.0001) or in combination with prazosin (t_357_ = -3.50, p = 0.0005) lowered sodium compared to placebo. Prazosin did not affect serum sodium. Females had lower minimum serum sodium than males (t_14_ = -2.43, p = 0.029). However, this did not appear to be a drug effect and was visible in their baseline measures. Eight of 448 serum sodium samples were not collected or not analyzable.

**Figure 3.**
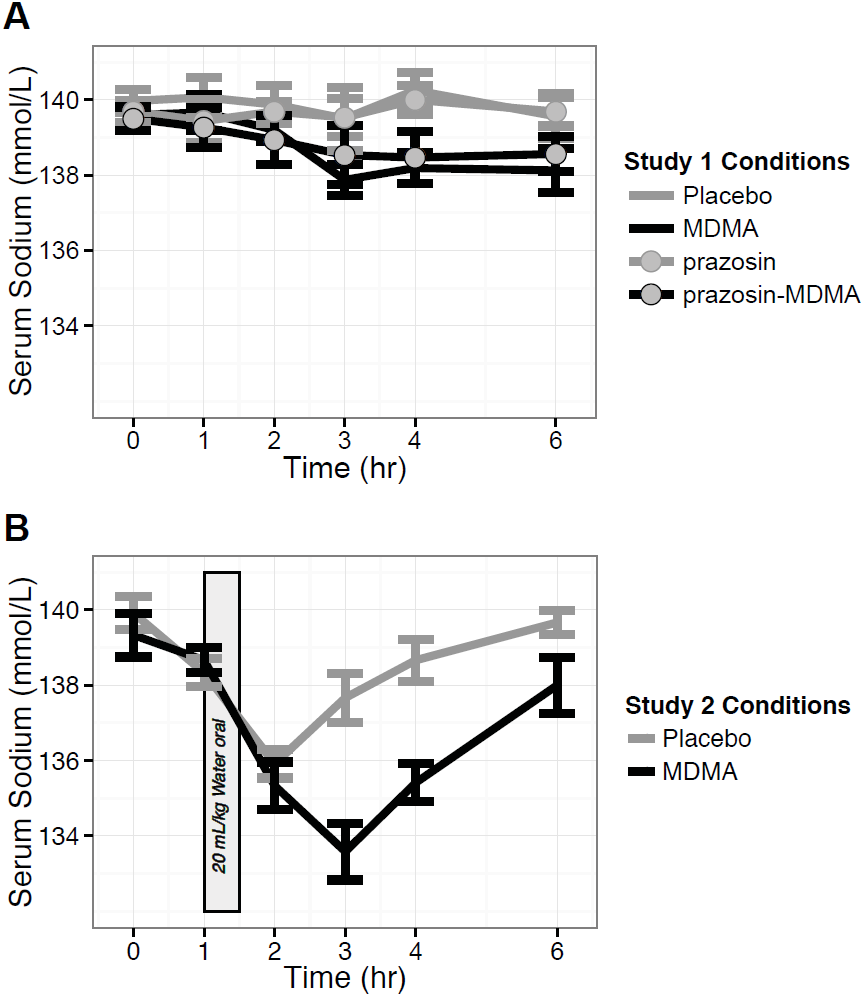
Serum sodium changes over time during Studies 1 (upper plot) and 2 (lower plot)

**Relationships between variables.** We did not detect any significant relationship between ADH and the decreases in serum sodium seen after MDMA. **Figure 4** provides a scatterplot of all ADH and serum sodium values collected after drug administration in the placebo and MDMA alone conditions. We saw no evidence that individual variability in pharmacokinetics influenced serum sodium: Peak MDMA and HMMA concentrations (and their interaction) also did not appear to predict significantly serum sodium decreases when included in a model that already contained drug condition.

**Study 2** included twelve (six male and six female) participants, ages 28.6± 1.9 years; range: 21-40). It differed from the first study in that participants in all conditions underwent oral water loading after an inpatient stay with standardized sodium and fluid intake. Females were tested during the follicular phase of their menstrual cycles. Two participants (one male, one female) vomited within 15 min of completing oral water loading during their MDMA conditions; these data are excluded from analysis. Study procedures were otherwise well-tolerated by all participants.

**Figure 4.**
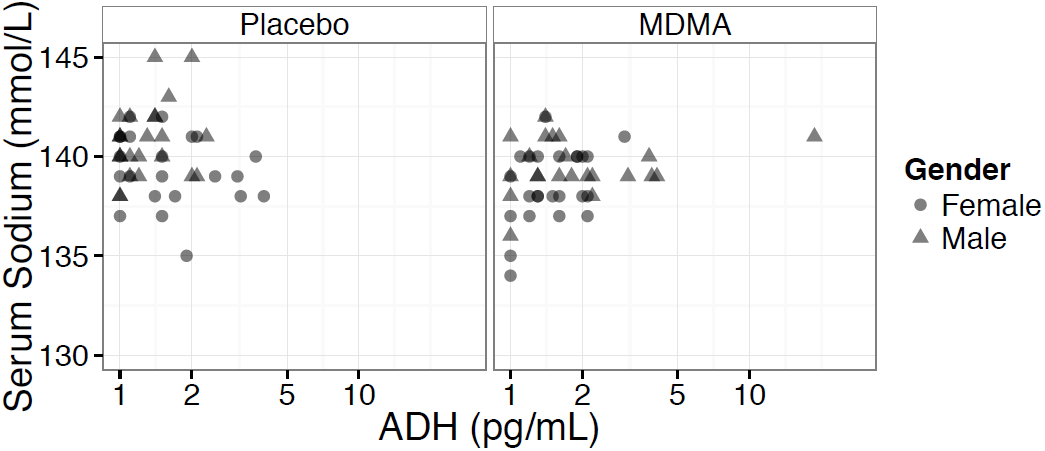
Relationships between Antidiuretic hormone (ADH) and serum sodium at 1, 2, and 4 h after Placebo and MDMA in Study 1.

**ADH and copeptin.** As in the previous study, we did not detect a significant effect of MDMA on ADH (Cmax: F_1,15_ = 0.113, p = 0.74). We also did not detect any effect of MDMA on copeptin (Cmax: F_1,9_ = 0.019, p = 0.89). Using data from all matching time points, there was a trend for ADH and copeptin to be correlated (Kendall's tau = 0.164, T=500, p = 0.068). Fourteen (8 from placebo) of the 120 ADH samples were not analyzable due to temperature control failures. One (from placebo, post-3-hr) of 72 copeptin samples was missing.

**Serum sodium.** MDMA with water loading decreased serum sodium to a greater extent than placebo with water loading (Cmin: F_1,9_ = 13.2, p = 0.005, **Figure 3, bottom**). Women tended to have lower baseline serum sodium, although their response to drug and water loading was not different from males. Specifically, including gender in the model revealed a trend for females to have lower values than males (Cmin: F_1,10_ = 3.93, p= 0.0755) without a significant interaction with condition. Copepetin also predicted serum sodium, apparently independently from condition (**Figure 5, right**).

In a model predicting serum sodium at 2 and 3 h after dosing, copepetin and condition each significantly predicted serum sodium (condition: F_1,32_ = 12.2, p = 0.001; copeptin: F_1,32_ = 4.22, p = 0.048), but there was no significant interaction. As visible in **Figure 5**, one female developed transient asymptomatic hyponatremia (serum sodium of 127 mmol/L) at 3 h after MDMA.

**Figure 5.**
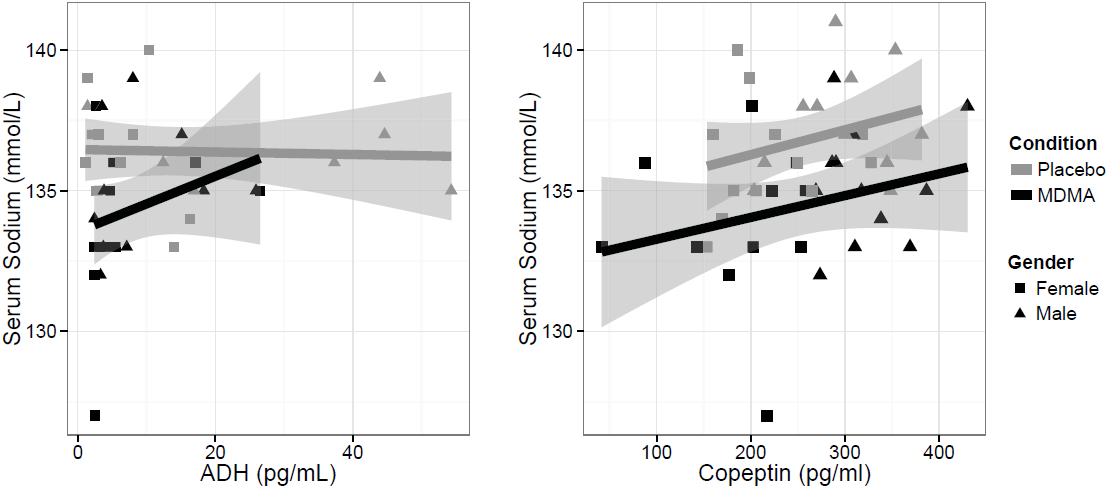
Relationships between Antidiuretic hormone (ADH, left), Copeptin (right), and serum sodium at 2 and 3 h after MDMA (black) or placebo (grey) in Study 2.

**Combined analysis.** In order to understand the interaction of MDMA and water intake, we analyzed serum sodium data from both studies combined (**Figure 6**). We estimated a model predicting serum sodium change from baseline using condition (i.e., the 4 combinations of drug and water loading) and time as fixed effects and participant as a random effect. This indicated there were significant effects of condition (F_3,278_ = 19.5, p < 0.0001) and time (F_5,278_ = 16.9, p < 0.001) and a significant interaction of the two (F_15,278_ = 8.62, p < 0.0001). We then tested each post-dose measurement time for the general linear null hypothesis that serum sodium after MDMA with water loading was significantly different from the sum of serum sodium changes after placebo water loading and serum sodium changes after MDMA with water restriction. This revealed significant nonadditive effects at 2 through 4 h (z = 3.56 to 5.19, p < 0.0005). In other words, MDMA and water loading have greater ability to lower serum sodium than either intervention alone.

**Figure 6.**
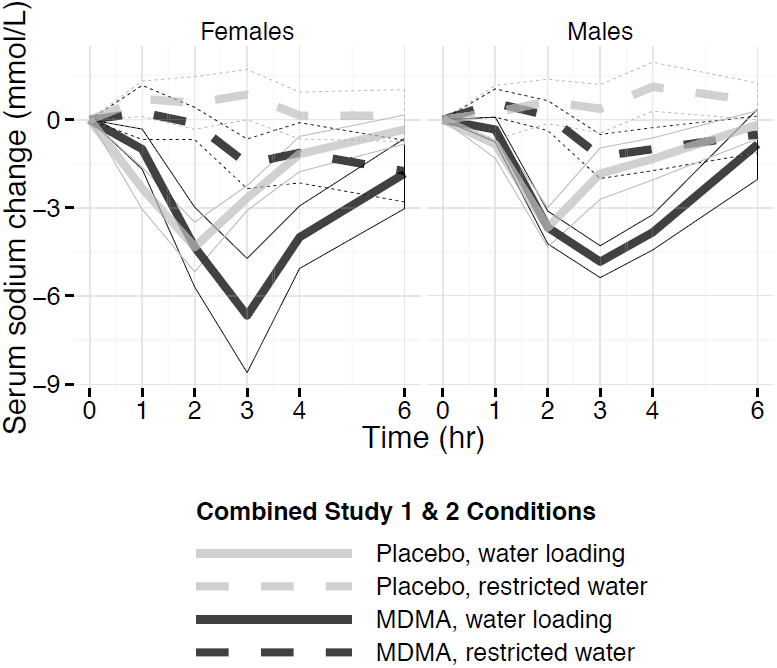
MDMA impairs serum sodium homeostasis after water loading. Plot shows effects of MDMA and water loading on serum sodium using data pooled across Studies 1 and 2.

## Discussion

Since its first description by Maxwell, Polkey, and Henry [5], there have been dozens of case reports of symptomatic hyponatremia in MDMA users, usually female [17, 54, 55]. Despite high rates of morbidity and mortality, there are few investigations of mechanisms of MDMA-related hyponatremia. The current manuscript describes the first controlled trial to evaluate the effect MDMA on water homeostasis in response to water loading. We find that MDMA can both lower serum sodium and exaggerate the hypnonatremic response to fluid intake in male and female participants. This suggests that symptomatic hyponatremia in MDMA users represents a severe form of a common drug effect rather than a rare idiosyncratic response. In addition, hypotonic fluid intake appears to be a risk factor for MDMA-related symptomatic hyponatremia.

Decrease in serum sodium after MDMA has been seen in two past clinical studies and is consistent clinical reports that MDMA increases risk of clinical hyponatremia. Two of three previous studies [21, 56] have detected a statistically significant decrease in serum sodium following MDMA administration compared with the placebo condition. Our studies support this phenomenon. In Study 1, both MDMA conditions (i.e., MDMA alone and MDMA with prazosin) resulted in a significant decrease in serum sodium. The one past report that did not detect change in serum sodium after MDMA did not control fluid intake beyond limiting total volume to 2 L and only made a single post-dose measure at 2 h [31]. In Study 2, the MDMA condition was associated with a larger decrease in serum sodium following an oral water challenge compared with placebo (and water loading). Pooling data from both studies, we found evidence that the combined effects of MDMA and water loading were greater than would be predicted from these manipulations measured individually.

We detected serum sodium changes after MDMA and water loading in both male and female participants. This may raise the question of why clinical cases of MDMA-related symptomatic hyponatremia are predominantly female. This may represent tendencies of females to ingest higher doses of MDMA and greater amounts of fluid compared to males when intake is normalized by body mass. However, females are also known to be at elevated risk of developing symptoms from hyponatremia of all causes [57, 58]. This is not thought to be due to a gender difference in probability of lowered serum sodium, but to a sex difference in ability of neurons to respond to lowered serum sodium. Estrogens in premenopausal females impair brain cell volume regulation by inhibiting Na*-K*-ATPase pumps that remove sodium from astrocytes. Moreover, ADH may potentiate cerebral vasoconstriction to a greater degree in females than males, decreasing oxygen delivery [40].

Our findings, if confirmed, suggest MDMA may cause what Zerbe, Stropes, and Robertson [59] refer to as Type C syndrome of inappropriate antidiuretic hormone secretion (SIADH). Type C SIADH manifests as failure to suppress ADH secretion at plasma osmolalities below the osmotic threshold. Plasma ADH shows a normal relationship with plasma at physiological plasma osmolalities, but ADH is inappropriately high at low plasma osmolalities. These relationships are suggested in Figure 5 in which copeptin release is affected by osmolality yet it is shifted in comparison to the control, while ADH release fails to suppress at serum sodium concentrations of less than 136 mmol/L.

The decreases in serum sodium we and other investigators detected in controlled MDMA administration studies are on average modest, with the exception of one case of asymptomatic hyponatremia after combined MDMA and water loading we detected in Study 2. It may be that the severe hyponatremia documented in case reports is due to a combination of factors, including excess intake of hypotonic fluids [19]. Polydipsia observed with MDMA use in case reports may be due to hyperpyrexia, a hypothesized change in primary drive to drink, and exposure to harm reduction messages emphasizing the need to avoid dehydration [19, 60]. Unfortunately, we did not measure thirst in either of our studies.

We were not successful at establishing a proximal mechanism of serum sodium decreases in MDMA users. Contrary to hypotheses we find no relationship between this lowered serum sodium and either ADH or the longer-lasting co-factor copeptin. Three previous studies [21, 31, 56] have reported an effect of MDMA on concentrations of ADH (or the more readily detectible co-factor copeptin), while we did not detect such an effect. In our first study, our positive control, prazosin, resulted in increased levels of ADH, while MDMA did not. In our second study, we also did not detect a significant increase in ADH or copeptin. It is possible that water loading suppressed drug-induced ADH and copeptin secretion in our second study (as was seen after infusion of desmopressin and hypotonic saline infusion in [30]). Variability in our ADH assay may have also made it difficult to detect drug effects. It is also worth noting that the ADH elevations reported in previous controlled MDMA studies were generally small in magnitude and did not appear dose dependent. Concentrations seen after 40 mg in males in the study of Henry et al. [21] are comparable to those reported after 125 and 100 mg MDMA in both genders in the papers of Simmler et al. [31] and Dumont et al. [25].

Our results raise the question of whether MDMA might impair water balance in part (or in whole) by a mechanism that is wholly or partly independent of elevated ADH. There are certainly other known mechanisms by which water balance can be impaired. Drugs such as carbamazepine and oxcarbazepine are thought to potentiate the effects of ADH at the renal tubule level rather than directly altering ADH levels [61]. Oxytocin, which is released by MDMA, is believed to regulate renal water reabsorption (reviewed in [62]), and should be examined in future studies.

Variations in MDMA metabolism are expected to contribute to variability in increases in ADH since HMMA may also contribute to ADH release [21–24] and genetic polymorphisms associated with low CYP2D6 or COMT enzyme activities were associated with greater MDMA-induced lowering of plasma sodium in a study of illicit ecstasy users [26]. However, we were not able to find significant relationships between either peak MDMA or HMMA concentrations and serum sodium decrease in our first study. Given the modest size of our study, these potential relationships deserve further investigation.

Our research supports the need for harm reduction measures to prevent complications from MDMA-related hyponatremia. Moritz, Kalantar-Zadeh, and Ayus [63] have proposed a protocol for this modeled after that from the Second International Exercise-Associated Hyponatremia Consensus Development Conference [64]. This includes having medical personnel present at large cultural events where drug use is common. These personnel should be equipped to do on-site analysis of sodium concentrations in individuals with symptoms suggesting of MDMA-related hyponatremia, such as nausea, vomiting, headache, confusion, lethargy, altered mental status or seizure. Moritz et al. suggest individuals presenting with MDMA-related hyponatremic encephalopathy be treated with one or more 100 mL bolus infusions of 3% NaCl to reduce brain edema [65]. Finally, MDMA users should be cautioned that overconsumption of hypotonic fluids represents a risk factor for hyponatremia, particularly in females [66].

Our research has several limitations. Both studies used single dose levels of study drugs. In our first outpatient study, participants’ hydration statuses may have varied at the start of the study. Our second inpatient study was designed to correct these limitations. Another limitation is that we did not measure self-reported thirst. It is important to establish to what extent MDMA may alter thirst or increase polydipsia [19]. Finally, we did not measure urine osmolality, which could have greatly aided interpretation of our results.

In conclusion, we found that MDMA lowers serum sodium but we were not able to relate this change to ADH or copeptin. We also found that MDMA acutely exaggerates the hyponatremic effects of water, indicating that hypotonic fluid intake is a risk factor for MDMA-related hyponatremia.

## Acknowledgment

This research was supported by National Institutes of Health DA017716 and DA016776 and the NIH/National Center for Research Resources UCSF-CTSI UL1 RR024131. The authors thank Ben Campbell, Ryne Didier, Margie Jang, Juan Carlos Lopez, and Jennifer Siegrist for assistance with participant recruitment, data collection, and regulatory compliance.

